# Arabidopsis late blight: Infection of a nonhost plant by *Albugo laibachii* enables full colonization by *Phytophthora infestans*

**DOI:** 10.1101/035006

**Authors:** Khaoula Belhaj, Liliana M. Cano, David C. Prince, Ariane Kemen, Kentaro Yoshida, Yasin F. Dagdas, Graham J. Etherington, Henk-jan Schoonbeek, H. Peter van Esse, Jonathan D.G. Jones, Sophien Kamoun, Sebastian Schornack

**Affiliations:** The Sainsbury Laboratory, Norwich Research Park, Norwich, United Kingdom.; Department of Plant Pathology, North Carolina State University, Raleigh, USA.; School of Biological Sciences, University of East Anglia, Norwich, United Kingdom.; Max Planck Institute for Plant Breeding Research, Cologne, Germany.; Organization of Advanced Science and Technology, Kobe University, Kobe, Hyogo, Japan.; The Genome Analysis Centre, Norwich Research Park, Norwich, United Kingdom.; Department of Crop Genetics, John Innes Centre, Norwich Research Park, Norwich, United Kingdom.; Sainsbury Laboratory, University of Cambridge, Cambridge, United Kingdom

## Abstract

The oomycete pathogen *Phytophthora infestans* causes potato late blight, and as a potato and tomato specialist pathogen, is seemingly poorly adapted to infect plants outside the Solanaceae. Here, we report the unexpected finding that *P. infestans* can infect *Arabidopsis thaliana* when another oomycete pathogen, *Albugo laibachii*, has colonized the host plant. The behaviour and speed of *P. infestans* infection in Arabidopsis pre-infected with *A. laibachii* resemble *P. infestans* infection of susceptible potato plants. Transcriptional profiling of *P. infestans* genes during infection revealed a significant overlap in the sets of secreted-protein genes that are induced in *P. infestans* upon colonisation of potato and susceptible Arabidopsis, suggesting major similarities in *P. infestans* gene expression dynamics on the two plant species. Furthermore, we found haustoria of *A. laibachii* and *P. infestans* within the same Arabidopsis cells. This Arabidopsis - *A. laibachii* - *P. infestans* tripartite interaction opens up various possibilities to dissect the molecular mechanisms of *P. infestans* infection and the processes occurring in co-infected Arabidopsis cells.

## Introduction

Plants have evolved diverse and effective mechanisms to protect against attack by microbial pathogens. Indeed, a central tenet of plant pathology is that resistance is the rule and disease the exception (Briggs, 1995). Although broad host-range pathogens do occur, most plant pathogens are adapted to a limited number of taxonomically related host species and cause disease on only a few host plants. Those pathogens may not fare well on plants unrelated to their hosts due to adaptive evolution, which tends to drive organisms towards specialization, for example through the accumulation of mutations that enhance virulence on one host but impair it on another (Tosa *et al*., 2006, Borhan *et al*., 2008, Ma *et al*., 2010, Raffaele *et al*., 2010, Dong *et al*., 2014, Dong *et al*., 2015). In addition, nonhost resistance and species-specific resistance serve to restrict the host range of plant pathogens (Schulze-Lefert *et al*., 2011, Senthil-Kumar *et al*., 2013, Lee *et al*., 2014). Physical barriers, such as fortified cell walls and a waxy cuticle, production of antimicrobial secondary metabolites, and cell-autonomous immunity all contribute to nonhost resistance (Fellbrich *et al*., 2002, Bettgenhaeuser *et al*., 2014, Miedes *et al*., 2014, Piasecka *et al*., 2015). Futher, cell autonomous immunity is multi-layered, involving pre-invasive defences as well as cell surface and cytoplasmic immune receptors that perceive pathogens (Dodds *et al*., 2010, Win *et al*., 2012). Thus, a pathogen’s ability to colonize a certain plant species includes its capacity to suppress or tolerate host immunity.

The oomycete plant pathogens include numerous host-specific species (Lamour *et al*., 2009, Thines *et al*., 2010, Fawke *et al*., 2015, Kamoun *et al*., 2015). These filamentous microorganisms are some of the most destructive plant pathogens and remain persistent threats to both farmed and native plants (Akrofi *et al*., 2015, Enzenbacher *et al*., 2015, Hansen, 2015, Roy, 2015). For example, the Irish potato famine pathogen *Phytophthora infestans*, the causal agent of late blight, recurrently endangers global food security (Fisher *et al*., 2012, Fry *et al*., 2015). *P. infestans* is thought to have a relatively narrow host range, infecting a few wild *Solanum* species in their native habitats of central Mexico and the Andes, as well as cultivated potato and tomato in most regions where these crops are grown (Grunwald *et al*., 2005, Fry *et al*., 2009, Goss *et al*., 2014). *P. infestans* can also infect other solanaceous plants, such as petunia and the experimental host *Nicotiana benthamiana* (Becktell *et al*., 2006, Chaparro-Garcia *et al*., 2011). However, this pathogen is not known to complete its full infection cycle on plants outside the Solanaceae. For example, the model plant *Arabidopsis thaliana*, a member of the Brassicaceae family, is fully resistant to *P. infestans* and is considered a nonhost (Vleeshouwers *et al*., 2000, Huitema *et al*., 2003, Lipka *et al*., 2005, Stein *et al*., 2006).

On Arabidopsis leaves, as on other nonhost plants such as tobacco and parsley, *P. infestans* cysts germinate, form appressoria, and directly penetrate epidermal cells to form infection vesicles and occasionally secondary hyphae (Colon *et al*., 1992, Schmelzer *et al*., 1995, Naton *et al*., 1996, Vleeshouwers *et al*., 2000, Huitema *et al*., 2003). However, this early interaction is followed by the hypersensitive response, a localized cell death reaction of plants that restricts the spread of the pathogen (Vleeshouwers *et al*., 2000, Huitema *et al*., 2003). In the Arabidopsis *pen2* mutant, which is deficient in the hydrolysis of 4-methoxyindol-3-ylmethylglucosinolate (4MO-I3M) into antimicrobial metabolites, the frequency of *P. infestans* penetration of epidermal cells increases, resulting in markedly enhanced hypersensitive cell death (Westphal *et al*., 2008). However, *P. infestans* does not complete its full infection cycle on *pen2* mutants or *pen2* mutants combined with mutations in other defense-related genes (Lipka *et al*., 2005, Westphal *et al*., 2008, Kopischke *et al*., 2013). In these mutants, *P. infestans* hyphae fail to extensively colonize the Arabidopsis mesophyll and do not develop haustoria, the specialized hyphal extensions that project into host cells and are thought to be sites where the pathogen secretes virulence proteins (effectors) (Whisson *et al*., 2007, Schornack *et al*., 2010). To date, there are no published reports of Arabidopsis mutants that are fully deficient in nonhost resistance to *P. infestans*, and thus enable extensive biotrophic colonization and sporulation of this pathogen (Stegmann *et al*., 2013, Geissler *et al*., 2015).

One oomycete pathogen that can infect *Arabidopsis thaliana* is *Albugo laibachii*, one of several specialist *Albugo* species that cause white blister rust disease (Kemen *et al*., 2011, Kamoun *et al*., 2015). *Albugo* spp. are obligate biotrophic parasites that are phylogenetically distinct from other oomycetes, such as *Phytophthora*, and thus have independently evolved the ability to colonize plants (Thines *et al*., 2010, Kemen *et al*., 2012). *Albugo* are widespread as endophytes in asymptomatic natural populations of Brassicaceae and likely influence the biology and ecology of their host species (Ploch *et al*., 2011). Remarkably, *Albugo* can suppress host immunity to enable colonization by other races of pathogens and subsequent genetic exchange between specialized genotypes with non-overlapping host ranges (McMullan *et al*., 2015). Prior infection by *Albugo* enhances susceptibility to plant pathogens such as downy and powdery mildews (Bains *et al*., 1985, Cooper *et al*., 2008). For instance, pre-infection with *A. laibachii* enables avirulent races of the Arabidopsis downy mildew *Hyaloperonospora arabidopsidis* to grow and sporulate on resistant Arabidopsis accessions (Cooper *et al*., 2008). *A. laibachii* suppresses the runaway cell death phenotype of the Arabidopsis *lesion simulating disease1* mutant, further supporting the view that this pathogen is an effective suppressor of plant immunity (Cooper *et al*., 2008). The mechanisms by which *Albugo* spp. suppress immunity remain unknown, but probably involve suites of effector genes like those identified in the *Albugo candida* and *Albugo laibachii* genomes (Kemen *et al*., 2011, Links *et al*., 2011).

Here, we aimed to determine the degree to which *A. laibachii* would enable maladapted pathogens to colonize Arabidopsis. Pre-infection with *A. laibachii* did not alter resistance of Arabidopsis to the Asian soybean rust pathogen *(Phakopsora pachyrhizi)* or the powdery mildew pathogen *(Blumeria graminis* f. sp. *hordei (Bgh))*. However, we discovered that pre-infection with *A. laibachii* enables the potato pathogen *P. infestans* to fully colonize and sporulate on Arabidopsis, a plant that is considered to be a nonhost of this Solanaceae specialist. Our results show that *P. infestans* carries the potential to infect other plant species outside its natural host spectrum employing a conserved set of transcriptionally induced effector genes. The interaction of Arabidopsis - *A. laibachii* - *P. infestans* will be an excellent model to examine how co-infection of host cells enables infection by *P. infestans*.

## Results

### *Albugo laibachii* infection enables *P. infestans* colonization of the nonhost plant Arabidopsis

Previous work indicated that the potato late blight pathogen *P. infestans* can penetrate epidermal cells of its nonhost Arabidopsis resulting in hypersensitive cell death but little ingress beyond the infection site (Vleeshouwers *et al*., 2000, Huitema *et al*., 2003). To determine the extent to which *A. laibachii* alters the interaction between Arabidopsis and *P. infestans*, we carried out serial inoculations with the two pathogens. First, we infected rosette leaves of 5-week-old Arabidopsis Col-0 with spores of *A. laibachii* strain Nc14 (Kemen *et al*., 2011). Successful infections were identified based on the formation of white sporangiophores on the abaxial side of rosette leaves 10 days after inoculation. At that stage, we detached the infected leaves, inoculated them with zoospores of *P. infestans* 88069 and monitored symptom development (Fig. 1A-B). Within 5 days after inoculation with *P. infestans*, we observed water-soaked tissue, necrosis, and ultimately sporulation in co-infected leaves (Fig. 1B). As controls we also applied *P. infestans* zoospores to uninfected leaves of *A. thaliana* Col-0, and also monitored mock- and *A. laibachii-inoculated* leaves (Fig. 1 A-B). No necrosis was observed in these negative controls (Fig. 1 A-B). To further investigate the degree to which *P. infestans* colonizes pre-infected Arabidopsis leaves, we repeated the experiment with *P. infestans* 88069td, a transgenic strain that expresses the cytoplasmic red fluorescent protein (RFP) marker tandem dimer, and monitored pathogen ingress by microscopy. This revealed an extensive network of hyphae in the co-infected leaves that extended to most of the leaf within just 3 days after *P. infestans* inoculation and sharply contrasted with the *P. infestans-only* treatment (Fig. 2). We also repeated the experiment with whole plants to ensure that the observed effect was not an artifact of the detached leaf assay. Here also, *P. infestans* triggered severe disease symptoms and formed an extensive hyphal network only in the mixed-infection leaves (Fig. S1).

**Figure 1.**
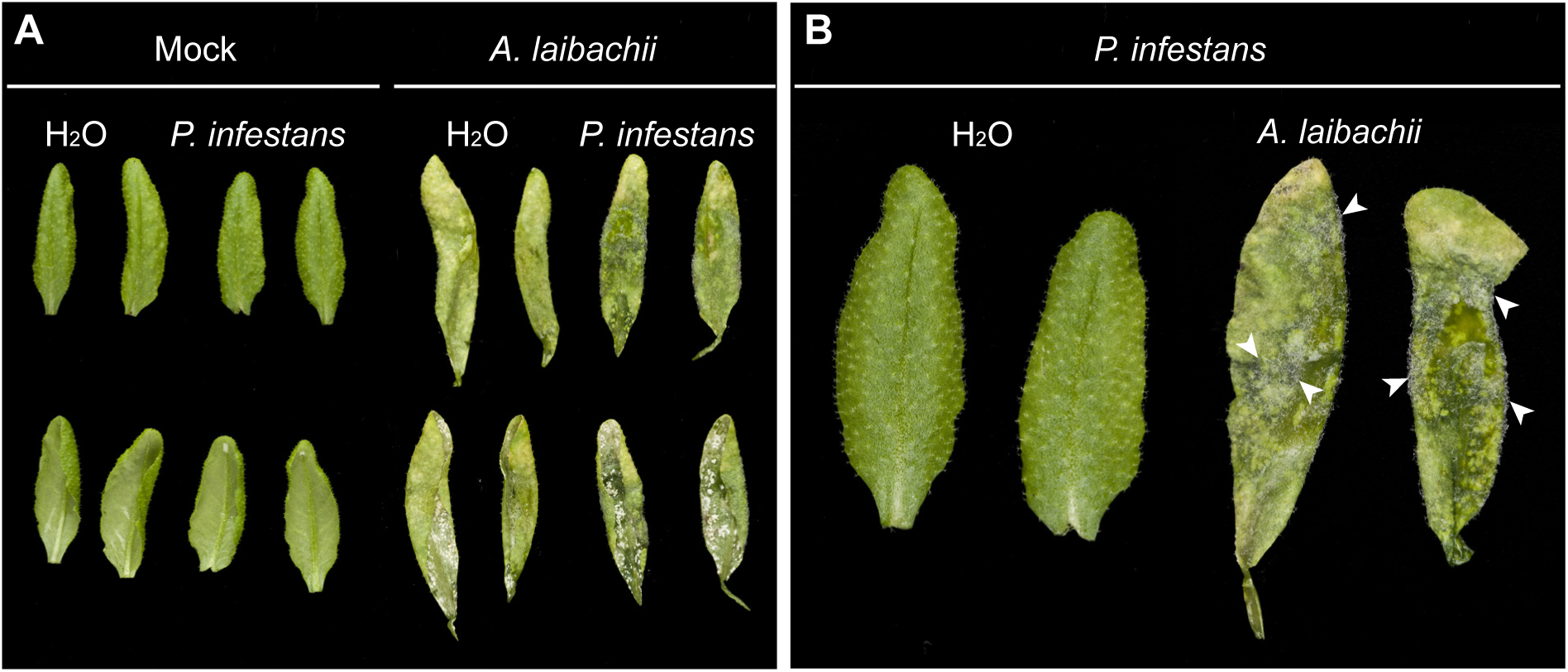
*Albugo laibachii* enables *Phytophthora infestans* to colonise Arabidopsis. (**A**) Control leaves (Mock) or leaves from *A. thaliana* Col-0 plants pre-infected with *A. laibachii* were detached and droplets of water (H_2_0) or *P. infestans* spore solution were applied to their abaxial sides and incubated for 4 days in high humidity. (**B**) A close up of (A) reveals *P. infestans* sporulation (arrowheads) as a dense cover of leaves pre-infected by *A. laibachii* only.

**Figure 2.**
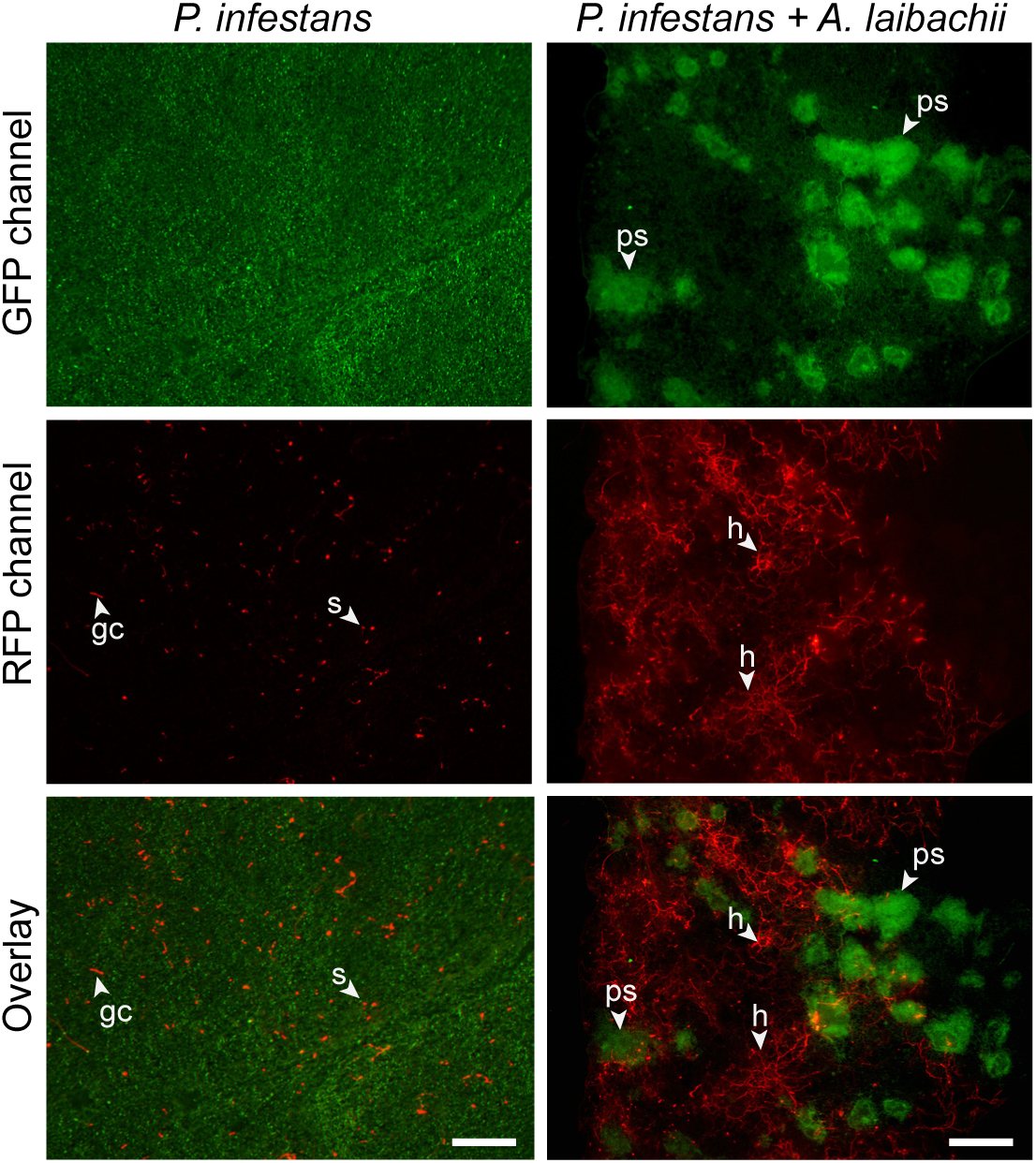
*A. laibachii* pre-infection supports extensive hyphal growth of *P. infestans* in Arabidopsis. Abaxial sides of control leaves of *A. thaliana* Col-0 (left column) and leaves pre-infected with *A. laibachii* (right column) have been infected with red fluorescent *P*. *infestans* 88069td. The extent of *A. laibachii* sporulation (visible as green accumulations under GFP illumination, upper row) and *P. infestans* hyphal colonisation (under RFP illumination, middle row) was assessed 3 days post inoculation using epifluorescence microscopy. Bottom row represents merged fluorescence pictures. Abbreviations: ps: pustules, h: hyphae; s: spores, gc: germinating cyst. Scale bar = 250 μm.

Next, we quantified pathogen biomass during infection using kinetic PCR as previously described (Judelson *et al*., 2000, Mauch *et al*., 2009) (Fig 3). We amplified the *P. infestans* gene *PiO8* to estimate relative levels of *P. infestans* DNA in infected plant tissue and observed a continuous increase over time in Arabidopsis leaves pre-infected with *A. laibachii* (Fig. 3). Overall, the pathology, microscopy, and molecular biology experiments indicate that *P. infestans* becomes able to fully colonize nonhost Arabidopsis plants upon pre-infection of those plants with *A. laibachii* (Fig. 1–3, Fig. S1).

**Figure 3.**
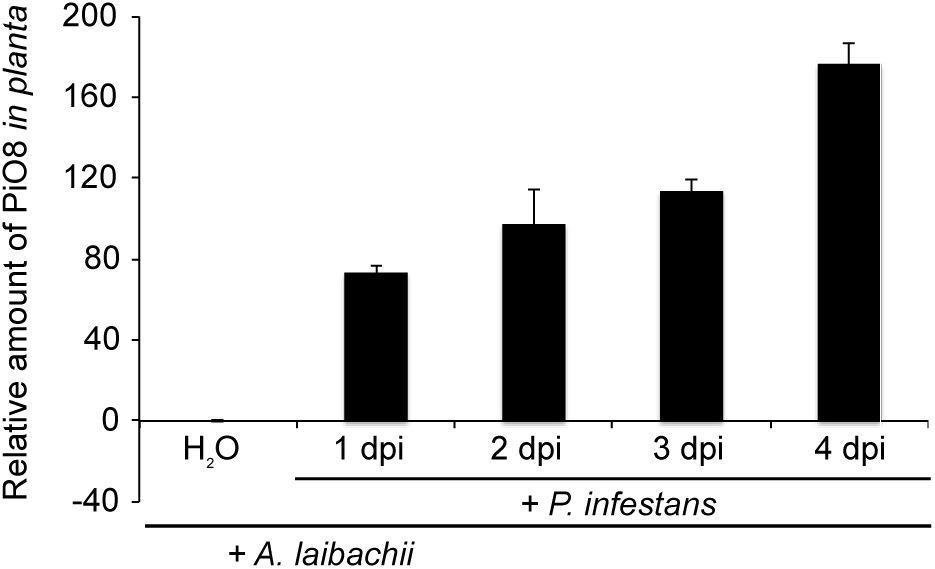
Quantification of *P. infestans* biomass upon infection of *A. thaliana* pre-infected with *A. laibachii*. Five-week old leaves of *A. thaliana* Col-0 pre-infected *with A. laibachii* were detached and drop-inoculated with *a* zoospore suspension of *P. infestans* isolate 06_3928A or mock-treated with water applied to their abaxial sides and incubated for 4 days under high humidity. DNA was extracted at 0, 1, 2, 3, and 4 days post inoculation and used for quantitative PCR (qPCR) for *PiO8* with gene-specific primers for *P. infestans*. DNA levels were normalized to the Arabidopsis *SAND* gene (At2g28390) and the relative amount of *Pi08* was normalized to the DNA level in mock-inoculated samples.

### Cellular dynamics of *P. infestans* colonization of pre-infected Arabidopsis

To study the interaction between *P. infestans* and pre-infected Arabidopsis in more detail, we performed confocal microscopy on leaves inoculated with *P. infestans* strain 88069td. We conducted side-by-side comparisons of the subcellular interactions of *P. infestans* in *A. laibachii-* and mock-infected leaves. In both cases, we observed germinated *P. infestans* cysts on the leaf surface as well as appressoria (Fig. 4A, Fig. 4B) and infection vesicles within the plant epidermal cells (Fig. 4C, Fig. 4D). The difference between the two treatments became apparent at 1 day post infection (dpi) with the activation of host cell death (the hypersensitive response, HR) at sites of attempted infection by *P. infestans* in mock-treated leaves only (Fig. 4E-F). Arabidopsis pre-infected with *A. laibachii* did not display an HR at sites of penetration of *P. infestans* (Fig 4G, Fig 4H). To independently validate these data, we inoculated *P. infestans* strain 88069td on leaves that were mock treated or pre-infected with *A. laibachii*, and then stained the leaves to quantify dead cells and monitor the invasion process at two different time points (6 hours post infection, hpi, and 24 hpi) (Fig. 4I). This again confirmed that penetration of *P. infestans* was not associated with the HR in samples that were pre-infected with *A. laibachii*, at both 6 and 24 hpi (Fig 4I).

**Figure 4.**
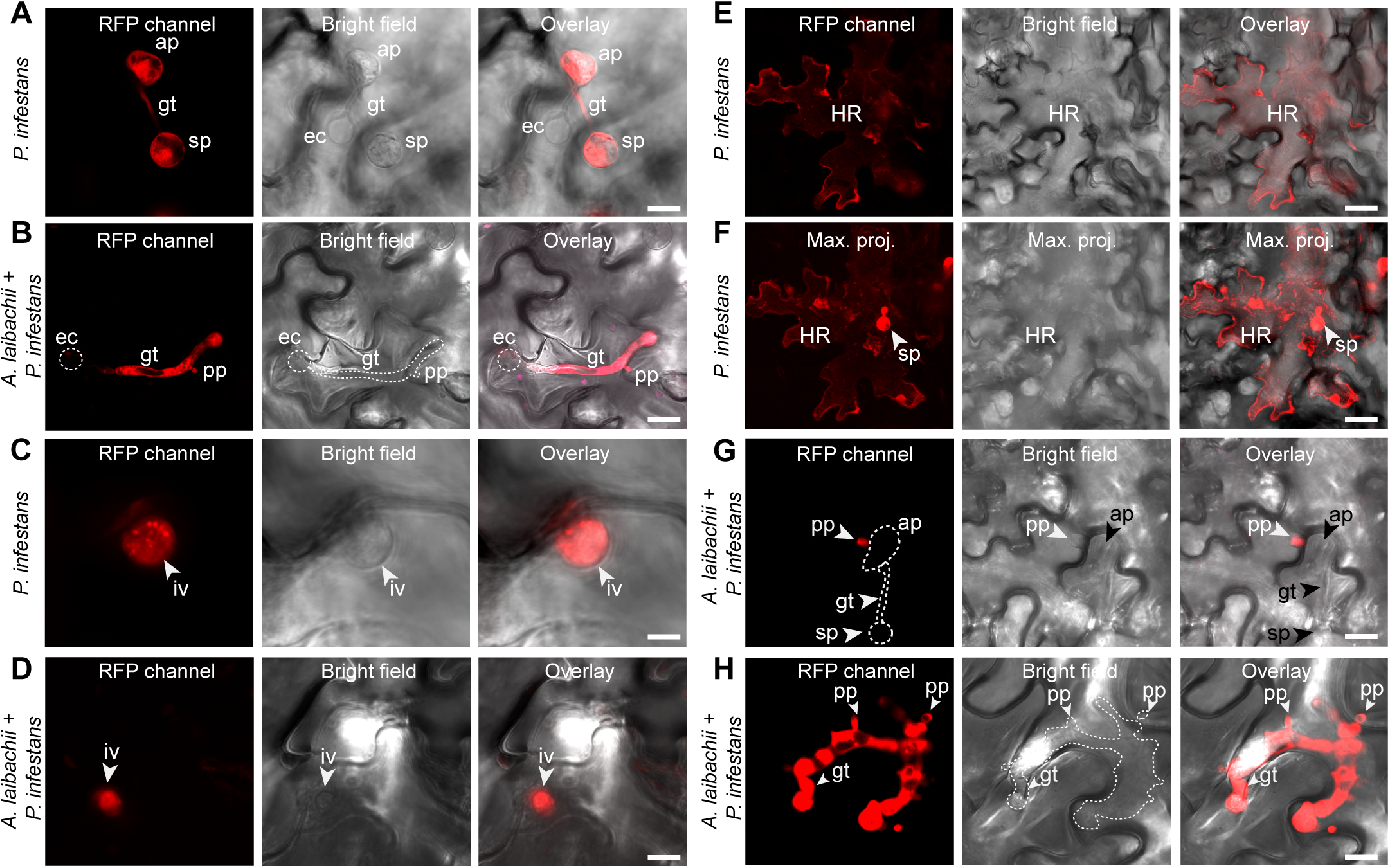
Hypersusceptibility of *A. thaliana* to *P. infestans* in leaves pre-infected with *A. laibachii* is accompanied by a loss of the hypersensitive response. Five-week-old leaves of *A. thaliana* Col-G mock-treated or pre-infected *with A. laibachii* were drop-inoculated with *a z*oospore suspension of red fluorescent *P. infestans* 88G69td Pathogen structures were visualized with confocal laser scanning microscopy at 16 hpi **(A-D)** and at 24 hpi **(E-H)** in samples treated with *P. infestans* only **(A, C, E, F)** and in co-infection experiments with *A. laibachii* **(B, D, G, H)**. Panel F represents a maximum projection of images produced from 18 Z stacks showing a hypersensitive response of the same area as panel E. **(I)** Counts of dead cells per leaf after infection with *P. infestans* in the presence or absence of pre-infection with *A. laibachii*. Data are representative of two biological replicates. Each replicate consists of counts from 8 independent leaves. Error bar represents ± SD. The two light microscopy inserts show examples of an HR cell death in infected leaves with *P. infestans* only (top panel) and of absence of HR cell death in co-infection experiments with *A. laibachii* and *P. infestans* (low panel) at 24 hpi. Abbreviations: sp: spores, gt: germ tube, ap: appressorium, ec: empty cyst, pp: penetration peg, iv: infection vesicle, HR: hypersensitive cell death, max. proj.: maximum projection. Scale bar = 25 μm (A-F) or 7.5 μm (G-H).

The ingress of *P. infestans* beyond its infection site became apparent starting at 36 hpi (1.5 dpi) in the *A. laibachii* pre-infected leaves, with intercellular hyphae spreading from the penetration site (Fig. S2). In contrast to mock-treated samples, the hyphae extended at 3 dpi to colonize the mesophyll and most of the leaf (Fig. S2). Branching hyphae with narrow, digit-like haustoria expanded from the site of penetration to neighboring cells through the intercellular space (Fig. S3). Starting at 3 dpi, the mycelium developed sporangiophores that released numerous sporangia to produce zoospores (Fig. S4). Thus, the *P. infestans* colonisation of Arabidopsis pre-infected with *A. laibachii* resembles, in behaviour and speed, the *P. infestans* infection reported on susceptible potatoes (Vleeshouwers *et al*., 2000).

### *A. laibachii* and *P. infestans* haustoria within a single Arabidopsis cell

*A. laibachii* forms haustoria in Arabidopsis cells (Caillaud *et al*., 2012). Since we observed the formation of haustoria by *P. infestans* in Arabidopsis pre-infected with *A. laibachii*, we searched for cells that harboured haustoria of both oomycetes. We recorded numerous events where single or multiple digit-like *P. infestans* haustoria co-occurred with multiple knob-like *A. laibachii* haustoria in the same Arabidopsis cells (Fig. 5). We found both haustoria types in epidermal as well as mesophyll cells of Arabidopsis. Thus, haustorium formation by *A. laibachii* or *P. infestans* does not trigger processes that prevent secondary penetration by another species. This observation will enable us to study how cell polarisation is affected by secondary penetration and how the two microbial pathogens vary in recruiting plant secretory processes to their haustoria.

**Figure 5.**
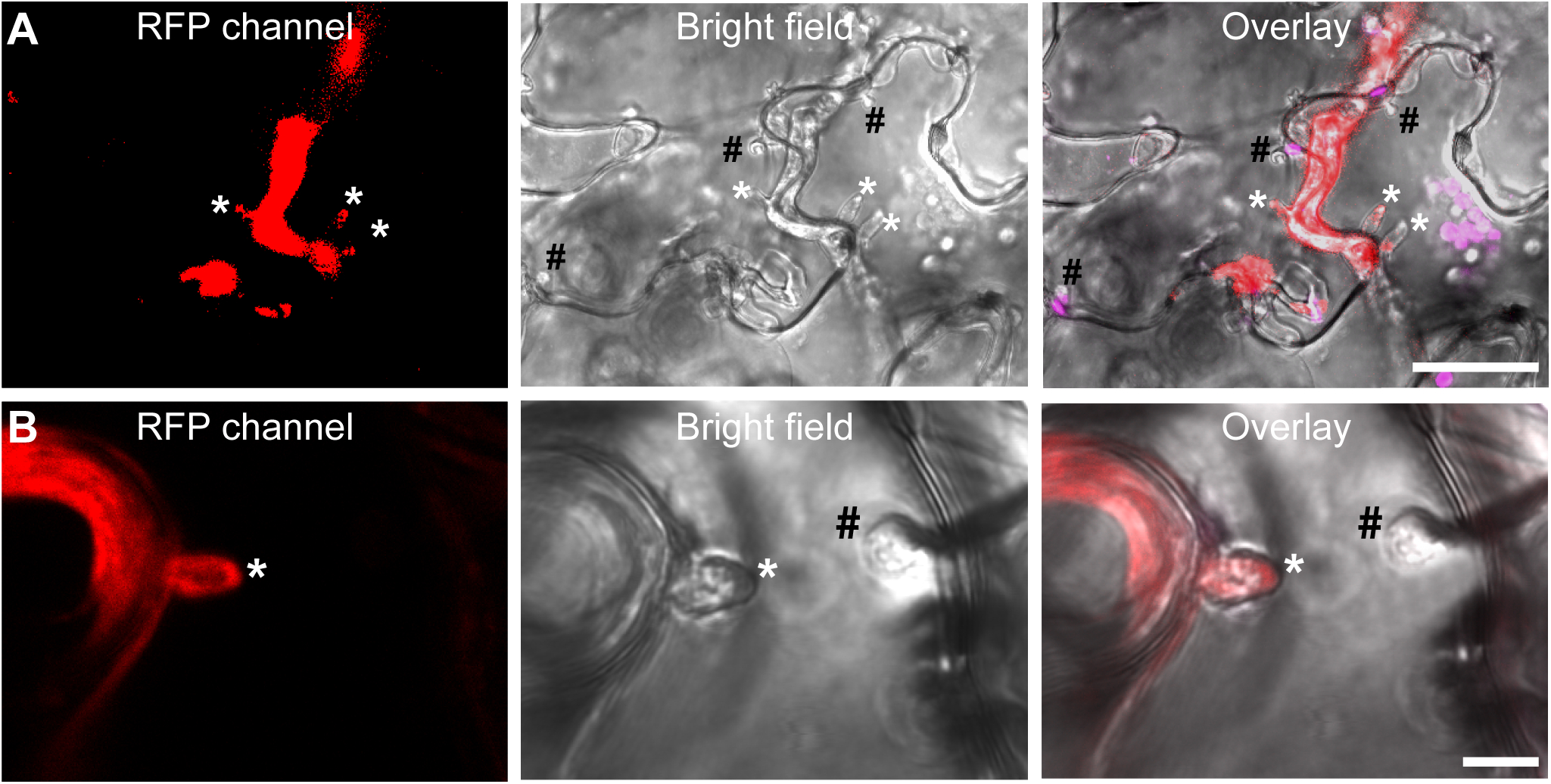
*Phytophthora infestans* and *Albugo laibachii* can form haustoria in the same Arabidopsis cell. (**A**) *A. thaliana* Col-0 precolonised with *A. laibachii* was infected with red fluorescent *P. infestans* 88069td. Inspection by microscopy at 2 dpi revealed the presence of haustoria. (**B**) Frequently, plant cells were observed to harbour digit-like, red fluorescent *P. infestans* 88069td haustoria as well as knob-like *A. laibachii* haustoria. Abbreviations: #: haustoria of *A. laibachii*, *: haustoria of *P. infestans*. Scale bar = 25 μm (A) or 7.5 μm (B).

### *In planta* expression dynamics of *P. infestans* secreted protein genes are similar on Arabidopsis and potato

Expression analyses have identified a significant set of *P. infestans* effector genes, which are transcriptionally induced during biotrophy in host-plant infections (Haas *et al*., 2009, Cooke *et al*., 2012, Pais *et al*., 2013). These studies have been limited to infections of potato and tomato, which both belong to the nightshade family (Solanaceae). To test whether the induced effector gene set is different in Arabidopsis pre-infected with *A. laibachii*, we collected *A. laibachii-infected* and mock-infected Arabidopsis leaves at different time points following application of zoospores of *P. infestans* strain 06_3928A (13_A2 clonal lineage, Cooke *et al*., 2012). To compare sets of differentially regulated *P. infestans* genes in Arabidopsis with those differentially regulated in potato, we also infected and harvested potato leaves. Extracted RNA from all samples was subjected to Illumina RNA-seq.

We found that during colonisation of potato, the steady-state transcript levels of 10,698 *P. infestans* genes were significantly altered. Of those, 7118 transcripts were also altered the same direction in *A. laibachii* pre-infected Arabidopsis. In contrast, 776 transcripts were exclusively altered in the *P. infestans* – *A. laibachii* – Arabidopsis interaction (see Supplementary Table 1 for details).

We next examined changes in transcripts encoding secreted proteins and found 196 induced sequences, of which 136 (66%) were shared, 40 were uniquely induced in Arabidopsis/A. *laibachii*, and 20 uniquely induced in potato (Fig S5, Fig. 6A). We found a strong correlation between Arabidopsis/A. *laibachii* and potato in the degree of gene expression induction both at 2 and 3 dpi with *P. infestans* (Fig.6B). Out of a total of 96 induced effector gene transcripts, a common set of 78 (81%) were induced in both plant species, whereas 12 and 6 effector transcripts where induced in a host-specific manner during colonisation of Arabidopsis and potato, respectively. Seven RXLR effector genes with known avirulence activity in specific potato cultivars were similarly induced in both host species (Fig. 6C). In summary, we conclude that the induction of secreted protein genes of *P. infestans* during colonisation of potato and Arabidopsis/A. *laibachii* leaves do not greatly differ.

**Figure 6.**
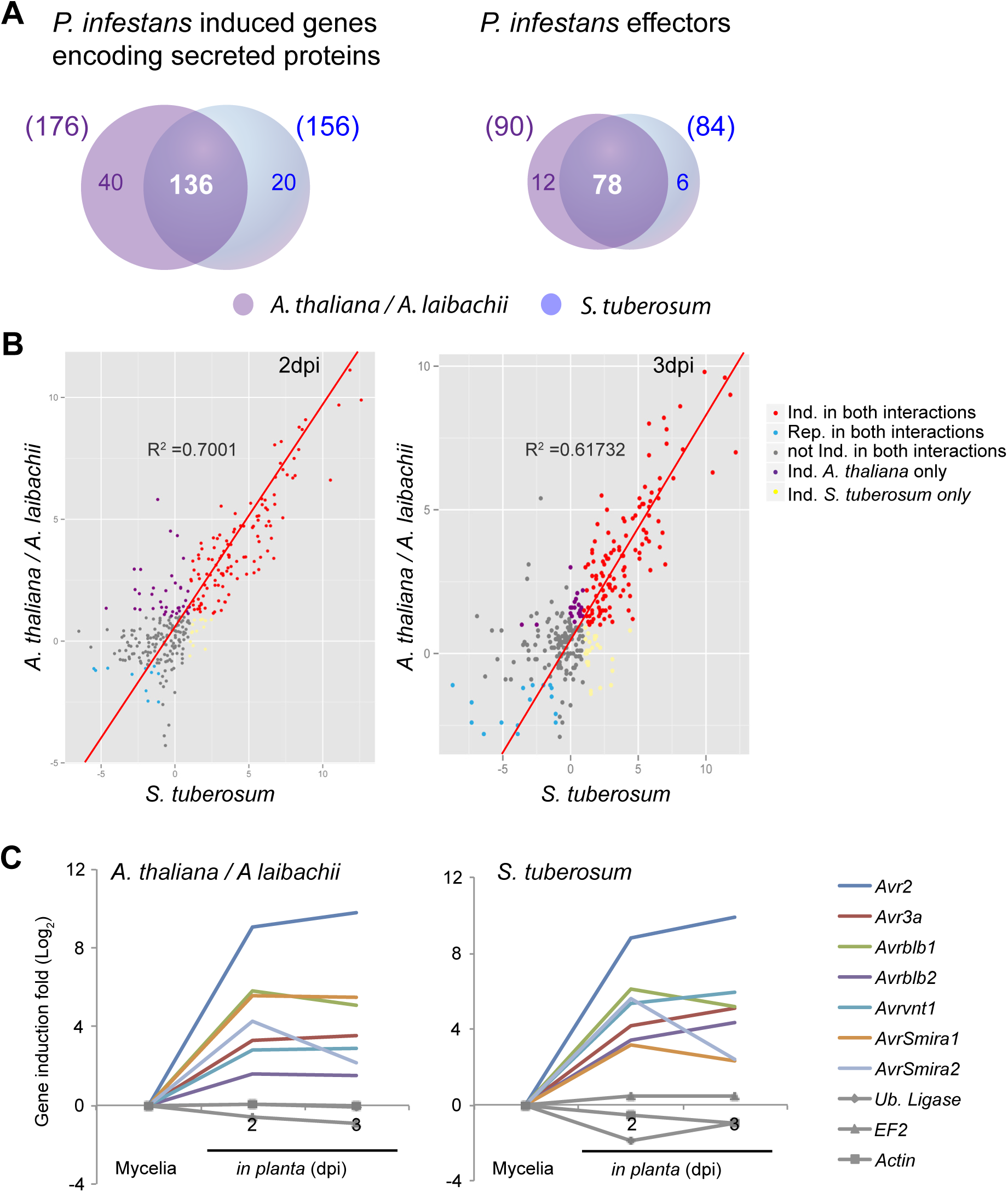
Similar sets of effectors are induced during *P. infestans* colonisation of potato *(Solanum tuberosum)* and Arabidopsis pre-infected with *A. laibachii*. (**A**) Numbers of commonly and uniquely induced genes encoding secreted *P. infestans* proteins and effectors (a subset of the secreted proteins). (**B**) Dot blot comparing the transcript levels of *P. infestans* effector-encoding genes between *S. tuberosum* and Arabidopsis pre-infected with *A. laibachii* at 2 (left panel) and 3 days post infection (right panel). (**C**) Gene expression intensities relative to the average expression intensity in media (Rye sucrose) are shown for genes encoding avirulence proteins during the interaction of *P. infestans* with *A. laibachii* pre-infected Arabidopsis leaves (left panel) and *S. tuberosum* leaves (right panel) induced at 2 dpi and 3 dpi. Genes encoding ubiquitin ligases, Elongation factor 2 and Actin are shown as uninduced controls. All expression intensities are log2 transformed.

### Arabidopsis leaves pre-infected with *Albugo laibachii* do not become susceptible to barley powdery mildew fungus or Asian soybean rust fungus

To determine the degree to which the effect of *A. laibachii* on *P. infestans* extends to other maladapted pathogens, Arabidopsis leaves pre-infected with *A. laibachii* were inoculated with the fungal pathogens *Blumeria graminis* f. sp. *hordei (Bgh)* and *Phakopsora pachyrhizi*, the agents of barley powdery mildew and Asian soybean rust, respectively. In both cases we observed no alteration of the interactions (Fig. S6 and S7). Both of these fungal pathogens failed to penetrate leaves of both mock- and *A. laibachii* pre-infected Arabidopsis plants.

## Discussion

In this study, we demonstrated that the potato blight pathogen *P. infestans* becomes capable of colonizing Arabidopsis when this nonhost plant is pre-infected by the obligate parasite *A. laibachii*. This is surprising, given that *P. infestans* is a Solanaceae specialist that is seemingly maladapted to plants from other botanical families. We took advantage of this tripartite interaction to perform comprehensive cellular and molecular analyses. On *A. laibachii-infected* Arabidopsis, *P. infestans* goes through its full infection cycle to a degree that has not been observed to date with pre- and post-invasive mutants (Lipka *et al*., 2005, Kobae *et al*., 2006, Stein *et al*., 2006, Westphal *et al*., 2008, Stegmann *et al*., 2013, Geissler *et al*., 2015). This includes the formation of haustoria, rapid hyphal proliferation, and profuse sporulation (Fig. 1–5, Fig. S1-S4). Expression dynamics of *P. infestans* genes encoding secreted proteins and effectors on susceptible (i.e., pre-infected) Arabidopsis were generally similar to those on potato, indicating that this pathogen colonizes pre-infected Arabidopsis in a similar manner as it colonizes its usual host plant (Fig. 6).

The finding that *P. infestans* can fully colonize immunosuppressed plants distantly related to its hosts indicates that pathogen host range may not be fully determined by a lack of essential factors in the nonhost, one of several resistance mechanisms generally thought to determine host specificity (Agrios, 2005). Indeed, there is little evidence that nonhost resistance results primarily from the absence of taxon-specific factors in the plant. For example, Garber’s nutritional theory, which postulates that resistant plants provide a “nutritional environment that is inadequate for a parasite” (Garber, 1956), has received little support over the years. By contrast, a greater understanding of the versatility and efficacy of the plant immune system has led to the view that active pre- and post-invasive defenses play a preponderant role in protecting most plants against most pathogens, and therefore in ultimately delimiting pathogen host range (Jones *et al*., 2006, Dodds *et al*., 2010).

Our findings are consistent with the evolutionary history of the *P. infestans* lineage, which reflects significant plasticity in host range. This lineage, also known as clade 1c, consists of a tightknit group of closely related species that have specialized on host plants from four different botanical families as a consequence of a series of host jumps (Grunwald *et al*., 2005, Raffaele *et al*., 2010, Dong *et al*., 2014). This indicates that on a macroevolutionary scale, the *P. infestans* lineage has the capacity to generate variants that can infect divergent host plants (Dong *et al*., 2015). The split between *P. infestans* and its sister species *P. mirabilis* is estimated to have occurred relatively recently ~1300 years ago (Yoshida *et al*., 2013), providing some indication of the frequency of host jumps within the clade 1c lineage.

*Albugo laibachii* converts Arabidopsis into a fully susceptible host of *P. infestans* to a degree that has not been observed to date with genetic mutants. The *pen2* and *pen3* mutants, which are deficient in penetration resistance, display enhanced responses to *P. infestans*, exhibiting a macroscopically visible hypersensitive cell death that results from increased frequency of epidermal cell penetration (Lipka *et al*., 2005, Kobae *et al*., 2006, Stein *et al*., 2006). However, the extent to which Arabidopsis penetration resistance to *P. infestans* is effective at stopping pathogen ingress is debatable given that penetration events can also be observed on wild-type Arabidopsis (Vleeshouwers *et al*., 2000, Huitema *et al*., 2003). In this study, we confirmed that penetration of Arabidopsis epidermal cells by *P. infestans* germinated cysts is commonly observed on wild-type Arabidopsis (Fig. 4). Thus, although *pen* mutants enable increased plant cell penetration, pre-invasive barriers do not fully block *P. infestans* infection, given that infection vesicles can be readily observed on mock-treated wild-type Arabidopsis at 16 hpi (Fig. 4). This view is consistent with the dramatic effect we observed on plants pre-infected with *A. laibachii*, which did not display *P. infestans-triggered* hypersensitivity probably as a consequence of post-invasive immunosuppression. Consistent with a post-invasive effect, *A. laibachii* did not alter Arabidopsis resistance to pathogens such as barley powdery mildew (Fig. S6) and Asian soybean rust fungi (Fig. S7), which cannot penetrate wild-type Arabidopsis cells, in sharp contrast to *P. infestans* (Fig. S2 and S3).

In *P. infestans*, as in many other filamentous pathogens, the expression of a subset of genes, notably secreted protein genes, is markedly induced during host infection (Haas *et al*., 2009, Cooke *et al*., 2012, Jupe *et al*., 2013, Pais *et al*., 2013). The mechanisms that underpin host signal perception by these pathogens, and the nature of these signals, remain largely unknown. We noted that the set of *P. infestans* effector genes induced on susceptible Arabidopsis largely overlaps with the genes induced in the host plant potato (Fig. 6). Patterns of effector gene expression displayed similar dynamics on both plants, with a peak during the biotrophic phase at 2 dpi. These results indicate that it is unlikely that *P. infestans* perceives a host-specific plant signal to trigger *in planta* gene induction. One possibility is that as the pathogen progresses from host cell penetration to intercellular hyphal growth to haustorium formation, it undergoes a developmental program that regulates gene expression.

Thines (2014) recently put forward the theory that *Albugo*-infected plants could serve as a bridge that enables other oomycetes to shift from one host plant to another. Indeed, repeated cycles of co-infection may facilitate the selection of genotypes of the maladapted pathogen that are virulent on the nonhost, eventually leading to a host jump. This scenario may have occurred with downy mildew species of the genus *Hyaloperonospora*, which tend to share Brassicaceae hosts with *Albugo* spp. (Thines, 2014). However, the degree to which *Albugo* has affected the ecological diversification of *P. infestans* and possibly other *Phytophthora* is unclear. First, it is not known whether the two pathogens are sympatric in central and south America, the natural geographic range of *P. infestans* and its sister species (Grunwald *et al*., 2005, Goss *et al*., 2014). Second, unlike *P. infestans*, most *Phytophthora* spp. are soil pathogens that do not spread aerially and are thus unlikely to colonize Albugo-infected leaves. Nonetheless, the possibility that biotic agents, such as *A. laibachii*, have facilitated host jumps in the *P. infestans* lineage should not be disregarded and deserve to be studied, for example by genome sequencing of environmental leaf samples. Our study further highlights the importance of studying multitrophic interactions in order to fully understand the biology and ecology of plant pathogens (Kemen, 2014).

Few diseases rival the effect of *P. infestans* on humankind (Fisher et al., 2012; Yoshida et al., 2013). Long after it triggered the Irish potato famine, this pathogen is still regarded as a threat to global food security and is an active subject of research (Kamoun et al., 2015). To date, *P. infestans* research has focused mainly on its interaction with Solanacaeous plants. Little progress has been achieved using model systems such as *Arabidopsis thaliana*, and work on Arabidopsis-P. *infestans* has been limited to studies of nonhost resistance (Huitema et al., 2003; Lipka et al., 2005; Kopischke et al., 2013). Other *Phytophthora* spp., e.g. *P. brassicae, P. cinnamomi, P. parasitica*, and *P. capsici*, have been shown to infect Arabidopsis but they have been hardly exploited in research (Roetschi et al., 2001; Robinson and Cahill, 2003; Belhaj et al., 2009; Wang et al., 2011; Wang et al., 2013). The Arabidopsis - *A. laibachii* - *P. infestans* tripartite interaction opens up several new avenues of research: 1) to address the genetic diversity of Arabidopsis resistance towards *P. infestans;* 2) to define the degree to which *Albugo* spp. have influenced the ecological diversification of *P. infestans* and enabled host jumps throughout evolution; 3) to dissect the molecular mechanisms, cell polarisation and retargeting of plant secretory pathways of co-infected host cells, a situation that is likely to occur frequently under natural conditions.

## Experimental procedures

### Biological material

*Arabidopsis thaliana* plants were grown on an “Arabidopsis mix” (600 L F2 compost, 100 L grit, 200g Intercept insecticide) in a controlled environment room (CER) with a 10 h day and a 14 h night photoperiod and at a constant temperature of 22°C. *A. thaliana* Col-0 ecotype was used for all experiments.

*P. infestans* isolate 88069 expressing a cytosolic tandem RFP protein (88069td) and *P. infestans* strain 06_3928A (13_A2 clonal lineage) were cultured on rye sucrose agar at 18°C in the dark as described earlier (Chaparro-Garcia *et al*., 2011, Cooke *et al*., 2012). *A. laibachii* strain Nc14 was used in pre-infection experiments in this study (Kemen *et al*., 2011). This strain was maintained on the *Arabidopsis thaliana* Col-5 line containing multiple insertions of the *RPW8* powdery mildew resistance gene (Col-gl *RPW8.1 RPW8.2)* (Xiao *et al*., 2001). The infected plants were kept overnight in a cold room (5°C) then transferred to a growth cabinet under 10-h light and 14-h dark cycles with a 21°C day and 14°C night temperature as described (Kemen *et al*., 2011). Besides *P. infestans* and *A. laibachii* we used two obligate fungal parasites: *Blumeria graminis* f.*hordei* CH4.8 (IPKBgh) and *Phakopsora pachyrhizi* isolate PPUFV02. A summary of fungal isolates used in this study and how they were maintained is provided in Supplementary Table 2.

### Sequential infection assays

All infection assays were performed on four- or five-week-old Arabidopsis plants of ecotype Col-0. Plants were pre-inoculated with a zoospore suspension of *A. laibachii* (7.5 × 10^5^ spores/ml) obtained from zoosporangia released from 14-day-old treated Col-gl *RPW8.1 RPW8.2* plants with *A. laibachii* isolate NC14 as described above. Briefly, whole Arabidopsis plants were sprayed with a zoospore suspension using a spray gun (1.25 ml/plant). They were incubated overnight in a cold room (5°C) in the dark and transferred later to a growth cabinet under 10-h light and 14-h dark cycles with a temperature of 21°C/14°C per day/night. Control plants were mock-treated with cold water. Plants pre-infected with *A. laibachii* NC14 were then used for second infections eight to ten days after inoculation with the pathogens listed in Supplementary Table 2. Co-infection assays with *P. infestans* were performed on detached leaves or whole plants as described earlier (Chaparro-Garcia *et al*., 2011). Briefly, a zoospore suspension of *P. infestans* (1 × 10^5^ spores ml^−1^) droplet was applied to the abaxial side of the leaf. Leaves were incubated on a wet paper towel in 100% relative humidity conditions with a 14 h/10 h day/night photoperiod and at a constant temperature of 18°C. Co-infection assays with powdery mildew pathogen (*Blumeria graminis* f. sp. *hordei* isolate CH 4.8) were performed on detached Arabidopsis leaves. Three-centimeter leaf strips were cut from the cotyledon or 1^st^ leaf of the barley cultivar and used as a control. Leaves were placed into agar plates containing 100 mg/l benzimidazole. Powdery mildew spores were collected from the barley-infected leaves on a piece of paper. Infection was made in a settling tower by tapping and blowing the inoculum. Plates were allowed to settle for 10 min after infection in the tower before incubation in a growth cabinet at 15°C (16 h light / 8 h dark with 18°C light / 13°C dark) (Brown *et al*., 1990). Co-infection assays with the Asian soybean rust were performed on detached leaves with *P. pachyrhizi* isolate PPUFV02 as described (Langenbach *et al*., 2013). Briefly, uredospores from *P. pachyrhizi-infected* soybean leaves were collected at 14 days-post inoculation (dpi), suspended in 0.01% (v/v) Tween-20 at 1 mg/ml and used for inoculation. Spore suspension of *P. pachyrhizi* was sprayed on Arabidopsis leaves until the droplets covered the whole leaf surface. To allow fungal spore germination, infected leaves were maintained in moist conditions (100% humidity) and in the dark for the first 24 hpi.

### Cytological analysis of infected material

Arabidopsis leaves infected with the red fluorescent *P. infestans* 88069td were visualized with a Fluorescent Stereo Microscope Leica M165 FC (Leica Microsystems Milton Keynes, UK) and an excitation wavelength for RFP: 510–560 nm. For confocal microscopy, patches of *A. thaliana* leaves were cut, mounted in water, and analyzed with a Leica DM6000B/TCS SP5 confocal microscope (Leica Microsystems) with the following excitation wavelength for the GFP and the RFP channels: 458 nm and 561 nm, respectively. Identical microscope settings were applied to all individuals. To quantify the HR cell death response in infected samples, leaves were stained with lactoglycerol-trypan blue and washed in chloral hydrate as described earlier (Belhaj *et al*., 2009). Specimens were mounted on microscope slides and analyzed with a Leica DM2700 M microscope (Leica Microsystems).

Powdery mildew structures were stained with lactoglycerol-trypan blue as described earlier (Vogel *et al*., 2000). Briefly, excised leaves were destained in ethanol overnight than washed thoroughly with in water and placed in lactoglycerol (1:1:1 lactic acid: glycerol: water). Specimens were mounted on microscope slides with a few drops of 0.1 % lactoglycerol-trypan blue staining on top. Fungal structures were imaged with a Leica DM2700 M microscope (Leica Microsystems).

Asian soybean rust-infected tissues were stained as described earlier in (Ayliffe *et al*., 2011). Briefly, Arabidopsis leaf tissue was placed in 1M KOH, then neutralized in 50 mM Tris, pH 7.0. The leaf was then stained with wheat germ agglutinin conjugated to fluorescein isothiocyanate (WGA-FITC, Sigma-Aldrich, UK) at 20 μg/ml. Specimens were mounted on a microscope slide and analyzed with a Leica DM6000B/TCS SP5 confocal microscope (Leica Microsystems) with an excitation wavelength for GFP of 458 nm.

All microscopy images acquired for the various infections were analysed by using the Leica LAS AF software, ImageJ (2.0) and Adobe PHOTOSHOP CS5 (12.0).

### Pathogen quantification

Genomic DNA was extracted from infected tissues using the DNeasy Plant Mini KIT (Qiagen, UK), following the manufacturer’s protocol. Quantification of pathogen growth *in planta* was performed by quantitative PCR using a rotor gene machine (Corbett Research Australia) as previously described (Mauch *et al*., 2009). The *PiO8* gene from *P. infestans* was used as a measure of *in planta* infection intensities of *P. infestans* with the following primers pair: PiO8-3-3F (5’-CAATTCGCCACCTTCTTCGA-3’) and PiO8-3-3R (5’-GCCTTCCTGCCCTCAAGAAC-3’) (Judelson *et al*., 2000). SYBR Green (Qiagen, UK) was used as fluorescent reporter dye to amplify the *PiO8* gene and was normalized to the Arabidopsis *SAND* gene (At2g28390) which was amplified with the following primer pairs SAND-F (5'-AACTCTATGCAGCATTTGATCCACT-3’) and SAND-R (5'-TGATTGCATATCTTTATCGCCATC-3’) (Mauch *et al*., 2009).

### RNA sequencing and analysis of the *P. infestans* transcriptome

We sequenced the following samples: i) 1 RNA sample from *P. infestans* isolate 06_3928A mycelia grown on RSA media, ii) 2 RNA samples from the dual interaction of *S. tuberosum* (potato cv. Desiree) infected with *P. infestans* isolate 06_3928A and iii) 3 RNA samples from the tripartite interaction of *A. thaliana* Col-0 sequentially infected with *Albugo laibachii* isolate NC14 and *P. infestans* isolate 06_3928A (Supplementary Table 3). These samples were labeled as: 1) *Phytophthora infestans* isolate 06_3928A mycelia grown on rye sucrose agar RSA (Pinf_mycRSA), 2) *P. infestans* isolate 06_3928A infecting *Solanum tuberosum* cv. Desiree and collected at 2 days post-incoculation (dpi) (Pinf_Stub_2dpi), 3) *P. infestans* isolate 06_3928A infecting *S. tuberosum* and collected at 3 dpi (Pinf_Stub_3dpi), 4) *Albugo laibachii* isolate NC14 colonising *Arabidopsis thaliana* Col-0 sequentially infected with *P. infestans* isolate 06_3928A and collected at 1 dpi (Alai_Atha_Pinf_1dpi), 5) *A. laibachii* isolate NC14 colonising *A. thaliana* sequentially infected with *P. infestans* isolate 06_3928A and collected at 2 dpi (Alai_Atha_Pinf_2dpi) and 6) *A. laibachii* isolate NC14 colonising *A. thaliana* sequentially infected with *P. infestans* isolate 06_3928A and collected at 3 dpi (Alai_Atha_Pinf_3dpi). Mycelium was harvested after being grown in liquid Plich media for 15 days. It was washed with distilled water, vacuum dried, and ground in liquid nitrogen for RNA extraction. Detached leaves of both plant species were inoculated with 10 μl of a zoospore solution of *P. infestans* isolate 06_3928A at 1 × 10^5^ spores ml^−1^. Leaf discs were collected at 2 and 3 days post inoculation (dpi) using a cork borer No. 4. Infected leaf samples were ground in liquid nitrogen until a fine powder was obtained and stored at −80°C prior to RNA extraction. We used the RNeasy Plant Mini Kit (Qiagen, Cat No. 74904), following the manufacturer’s instructions, to extract total RNA for all samples. cDNA libraries were prepared from total RNA using the TruSeq RNA sample prep kit v2 (Cat No. RS-122-2001). Library quality was confirmed before sequencing using the Agilent 2100 Bioanalyzer (Agilent Technologies). Sequencing was carried out using an Illumina Genome Analyzer II (Illumina Inc) with TruSeq Cluster generation kit v5 (Cat No. FC-104-5001) and TruSeq Sequencing kit v5 (Cat No. PE-203-5001). We performed read quality control by removing reads containing Ns and reads with abnormal read length (other than 76 bases) using FASTX-Toolkit version 0.0.13 (http://hannonlab.cshl.edu/fastx_toolkit). Total reads (76 bp paired-end) that that passed the parameters mentioned above for quality control were used for downstream analyses (Supplementary Table 3).

To extract read expression data of *P. infestans* isolate 06_3928A from the infected samples, we aligned each experiment to the genome assembly of *Phytophthora infestans* T30-4 version 2_2 (Haas *et al*., 2009) using TopHat software package version 2.0.6 (Kim *et al*., 2013) with 200 bp as the insertion length parameter. The alignments we obtained in sam format from TopHat software (Kim *et al*., 2013) were used for gene expression analysis (Supplementary Table 3). A two-stage analysis of the pathogen reads was applied to rescue multi-mapped or ambiguous reads that cannot be uniquely assigned to groups of genes. First, we generated Reads Per Kilo Base per Million (RPKM) values for each gene by using the htseq-count script that is part of the HTSeq python module (Anders *et al*., 2014). Next, we rescued reads that were enriched for gene families using multi-map group (MMG) approach and customized perl scripts (Robert *et al*., 2015). In brief, we allocated multi-mapped reads based on probability of multi-mapped reads derived from particular locus that was calculated from RPKM, and then estimated final RPKM according to a published method (Mortazavi *et al*., 2008). The adjusted-RPKM values of all reads after rescues were transformed into Log2 fold values by dividing the RPKM data to the RPKM values from mycelium of *P. infestans* isolate 06_3928A (Wagner *et al*., 2012). *In planta*-induced genes exhibiting at least two-fold gene induction between averaged media and infected sample (at 2 and/or 3 dpi) were considered induced during infection. Log2 values were loaded in Mev4_8 version 10.2 TM4 microarray software suite (Saeed *et al*., 2003) and analysed using hierarchical clustering method, gene tree selection, average linkage method and Pearson correlation for distance metric selection. The gene expression heatmap obtained with Mev4_8 shows fold-induction for *P. infestans* (PITG) genes with gene descriptions that are color-coded and highlight effector type custom annotations (supplementary figure 5).

## Acknowledgements

We thank Stephen Whisson for providing the *P. infestans* 88069td strain, Francesca Stefanato for providing *B. graminis* f. sp. hordei CH4.8, Oliver Furzer and Wiebke Apel for supplying plant material, and Matthew Moscow for providing WGA-FITC. This work was supported by the Gatsby Charitable Foundation, the European Research Council (ERC) and the Biotechnology and Biological Sciences Research Council (BBSRC). KY was supported by Japan Society of the Promotion of Science. H.S. was supported by the Institute Strategic Programme on Biotic Interactions for Crop Productivity BB/G042060/1.

## Author contributions

K.B., S.S. and S.K. designed experiments. K.B., L.M.C., D.C.P., H.S. and H.P.v.E carried out experiments. K.B, L.M.C., K.Y., G.J.E. and Y.F.D. analyzed data. D.C.P, A.K., H.S. and H. P.v.E. provided materials. K.B. S.K. and S.S. wrote the manuscript. S.K. and S.S. contributed equally to this work.

